# Urbanization plays a minor role in the flooding and surface water chemistry of Puerto Rico’s mangroves

**DOI:** 10.1101/423434

**Authors:** Benjamin L. Branoff

**Author notes:** ORCID 0000-0002-8796-2039 phone: 386 − 506 -7997 fax: 787 − 764 - 2610.

## Abstract

Although hydrology and water chemistry are known to change in proximity to cities, there remains little empirical evidence connecting specific components of urban landscapes to mangrove flooding dynamics or surface water chemistry. This study constructs five-year water level models from tidal harmonics and precipitation inputs to characterize mangrove flooding across urban gradients in three watersheds of Puerto Rico. There was some evidence for an influence of urbanization on both flooding and water chemistry, but this depended on the definition of urbanness, and points instead to geomorphology as the primary culprit. Urban sites exhibited 46% longer hydroperiods and 450% lower depths than non-urban sites. Rainfall importance was explained more by geomorphology than by urbanization and suggested systems with limited tidal connectivity are four times more sensitive to rainfall than systems with full tidal connectivity. There was also evidence for changes in tidal amplitudes along the urban gradient, which may explain the observed differences in flooding. Relationships between surface water chemical metrics and land cover contradicted previous studies by suggesting lower nutrients and biochemical oxygen demand with increasing urbanization. These results reinforce the understanding that the most important drivers of urban mangrove hydrology and water quality in Puerto Rico are likely geomorphology and tidal connectivity, with little but not zero influence from surrounding land cover. Results should be considered alongside the reported errors stemming from inaccuracies in digital elevation and rainfall response models, and will be useful in understanding future ecological censuses on the island.

## INTRODUCTION

As forested tidal wetlands along primarily tropical and subtropical coastlines, mangroves are often under pressure from urban development and the subsequent effects on surface water chemistry and hydrology, both of which play important roles in ecosystem function and the provisioning of services (Lugo and Snedaker 1974; Wolanski et al. 1993; Ewel et al. 1998; Medina 1999; Bosire et al. 2008; Lee et al. 2014). Specifically, metrics of flooding frequency, duration, and depth have been singled out as most important in influencing mangrove physiology and zonation (Krauss et al. 2006; Bosire et al. 2008; Lugo and Medina 2014). Likewise, concentrations of nitrogen and phosphorus as well as salinity and temperature have also been shown to be important indicators of mangrove nutrition, stress, and function (Medina 1999; Feller et al. 2003; Lovelock et al. 2009). Tides and rainfall contribute to system hydrology and water chemistry (Twilley and Chen 1998; Sutula et al. 2001), but groundwater, fluvial, and anthropogenic contributions may also be important in some systems (Ellison and Farnsworth 1996; Lee et al. 2006; Sakho et al. 2011; Gleeson et al. 2013). Thus, variations in geomorphology, which influence the breadth of the above hydrological contributions, is another important consideration when assessing mangrove ecosystems (Kjerfve et al. 1999; Adame et al. 2010). The complexity of this suite of potential influences is likely exacerbated in highly urban systems, where surface water flow and chemistry are directly and indirectly altered through impervious surfaces, infrastructure, and engineering projects (Leopold 1968; Hollis 1975; McClelland and Valiela 1998; Lee et al. 2006; Dietz and Clausen 2008). Although urban mangrove hydrology has been shown to be abnormal in some cases, few studies identify or quantify the specific influence of urbanization on hydrology or surface water chemistry.

Lee (2006) argues that the primary influence of urbanization on coastal wetlands is either directly or indirectly a result of changes to hydrology and sedimentation, which vary spatially and temporally in intensity. This is corroborated for forested wetlands in general (Faulkner 2004), but mangroves are unique in their tidal connectivity and specific examples from urban systems are scarce. The influence of municipal sewage discharge on mangroves has been well documented and generally leads to nutrient enrichment, heavy metal contamination, and sometimes mortality (Clough et al. 1983; Mandura 1997; Wong et al. 1997; Branoff 2017). Further, a number of studies point to changes in mangrove coverage following large engineering projects that alter surrounding geomorphology, but these are not necessarily related to urbanization, or use qualitative or otherwise ambiguous definitions of “urban” (Colonnello and Medina 1998; Tian-Hong et al. 2008; Sakho et al. 2011; Marois and Mitsch 2017). Studies directly linking quantified variations in urbanization metrics, such as impervious surface coverage, population density, or forested area, with changes in mangrove flooding dynamics or surface water chemistry are absent from the literature.

In San Juan, Puerto Rico, the mangroves have long been associated with ecological abnormalities due to ongoing urbanization (Seguinot Barbosa 1996), but again, there remains no direct ties between the relative intensity of urbanization and changes in flooding metrics or surface water chemistry. The geomorphology of the San Juan Bay Estuary has been modified through canalization or dredging or both over the course of the city’s 500 year history (Ellis 1976; Cerco et al. 2003). This has resulted in changes in underlying aquifer levels and salinities, as well as surface water chemistry and connectivity that are often associated with risks to both human and ecological health (Seguinot Barbosa 1996; Bunch et al. 2000). Large-scale engineering projects have since been proposed to mitigate these issues (Cerco et al. 2003), and are currently in various phases of implementation. However, apart from sewage discharge, there has been no reported effort to understand how specific components of the urban landscape (e.g. impervious surfaces, roads, population density) may be influencing mangrove flooding metrics or surface water chemistry. This holds true for the various other coastal urban watersheds of Puerto Rico, where mangroves have endured a long history of anthropogenic influences (Martinuzzi et al. 2009).

This study aims to characterize and model surface water fluctuations in the mangroves of three watersheds in Puerto Rico, using both long and short-term water level recordings, as well as rainfall and tidal harmonics models. It then uses these models alongside digital elevation models to analyze the flooding dynamics of the mangroves and correlates these, as well as surface water chemical properties, with surrounding land cover along gradients of urbanization. Specifically, this study tests the hypothesis that increasing surrounding urbanization is associated with variations in mangrove flooding and surface water chemistry. These connections must be well-understood for successful mangrove management (Lewis 2005; Bosire et al. 2008), and will be important in understanding the influence of urban land cover on mangrove ecology, especially in the context of future floral and faunal censuses in the study areas.

## METHODOLOGY

Three watersheds in Puerto Rico were selected based on the range in urbanization surrounding their mangroves. This was calculated as described below using spatial datasets of urban, open water, and vegetation classifications, as well as mangrove extent, population density and road lengths. These were combined as an urban index value to give a single relative representation of urbanization for one hundred random mangrove locations in all watersheds of Puerto Rico (Figure 1a & 1b). Sites were then selected within three watersheds to give the greatest possible range in urbanization within the mangroves (Figure 1c).

**Figure 1.**
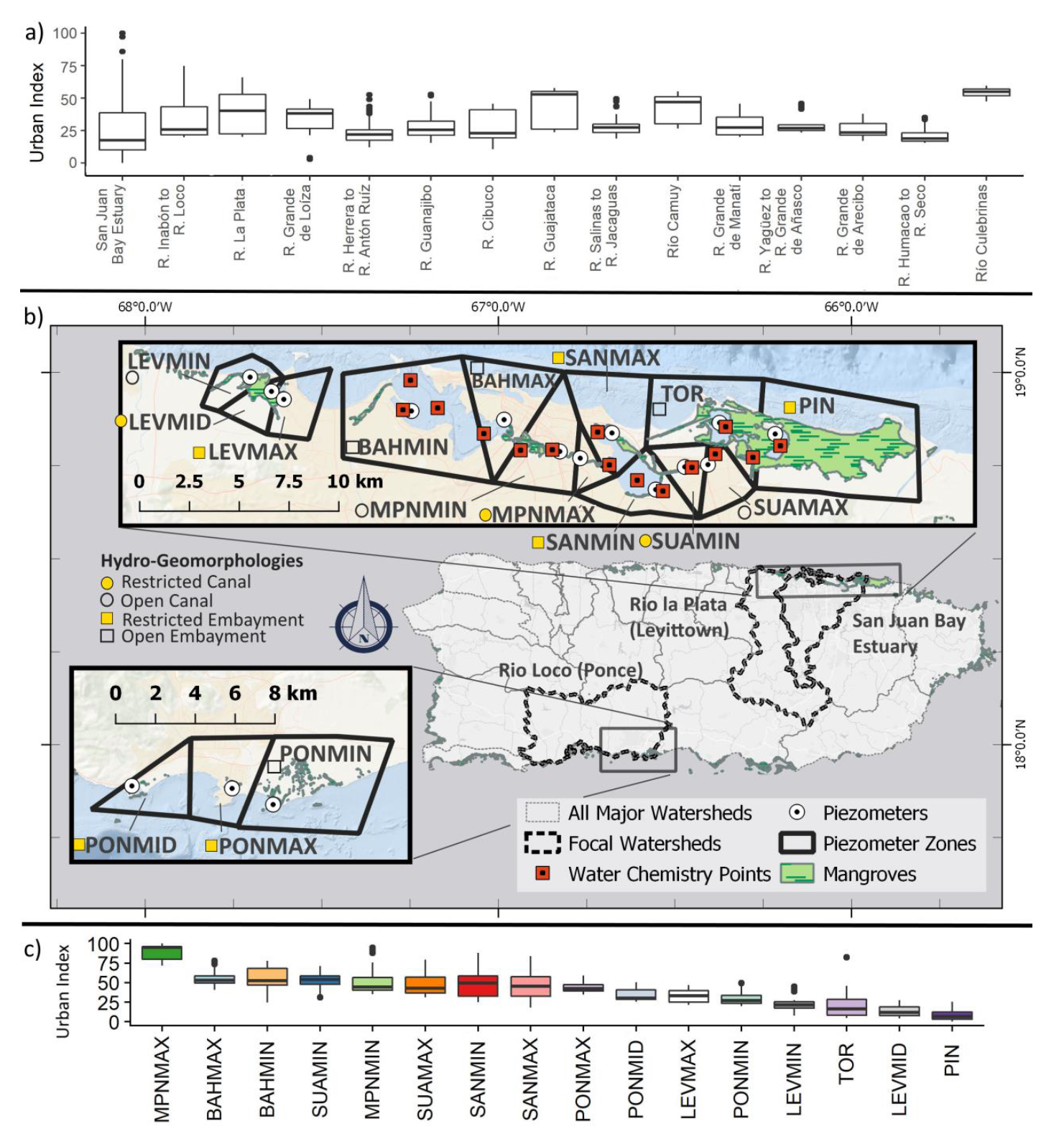
Study sites were selected by first choosing the three watersheds with the greatest range in urban index (a, ordered form left to right in decreasing urbanization range). The urban index is a combination of urban land use, vegetation and open water coverage, population density, and road length. Piezometers were placed within these watersheds (b), resulting in a gradient of urbanization across all sites (c). Inlay polygons represent individual piezometer zones. Water chemistry measurements were only available for the San Juan Bay Estuary. “MAX” and “MIN” postscripts refer to urbanization levels within each zone. BAH is the San Juan Bay, MPN is the Caño Martín Peña, SAN is the San José lagoon, SUA is the Suarez canal, TOR is the Torrecillas lagoon, PIN is the Piñones lagoon, LEV is Levittown and PON is Ponce.

### Study Locations

Three watersheds were chosen because their mangroves held the greatest range of urbanization. They are the San Juan Bay Estuary, the Río Inabón to the Río Loco (Ponce), and the Río la Plata (Levittown) (Figure 1a). The San Juan Bay estuary was divided into ten separate mangrove regions, and Ponce and Levittown into three regions each. This resulted in sixteen study locations throughout the three watersheds.

The San Juan metropolitan area lies within the watershed described as between the Río Grande de Loíza and the Río Bayamon. The 240 km^2^ drainage basin encompasses 25 km^2^ of open water, consisting of five embayments connected by a system of natural and canalized channels (Bunch et al. 2000). The estuary is home to the island’s largest and densest human population, 0.5 million people and 1,800 people/km^2^ in the municipalities of San Juan, Carolina and Bayamón (United States Census Bureau 2016). Given the intense human presence around the estuary, a number of studies have described its hydrodynamic and contamination issues, which range from eutrophication and heavy metals to tidal flushing and connectivity (Webb and Gómez-Gómez 1998; Bunch et al. 2000; Cerco et al. 2003; Acevedo-Figueroa et al. 2006). The estuary also harbors the islands largest mangrove forest, Bosque Estatal de Piñones, which at 2,550 ha represents roughly 11% of the watershed area (Brandeis et al. 2014). The combination of intense urbanization and expansive forested and open water areas within this watershed results in the greatest range in mangrove urbanization on the island. The median annual rainfall for this site was 1,463 mm from 2012 to 2017 (this study).

Dredging and canalization have occurred throughout the estuary and continue to be proposed (Ellis 1976; Seguinot-Barbosa 1983; United States Army Corps of Engineers 2015). Previous dredging has taken place in the San Juan Bay, and the Condado, San Jose and Torrecillas lagoons from the 1950s to the 1970s (Ellis 1976). These projects increased lagoon volumes and flow rates through canals, and decreased surface areas where dredging material was placed along shorelines. It is expected that this has resulted in lower residence times for sewage effluents and storm-water discharge, but greater prevalence of anaerobic zones.

Levittown refers to the mangroves associated with the town of Levittown, in Toa Baja, Puerto Rico, which falls within the Río la Plata watershed, just west of the San Juan Bay Estuary. The estuary is composed of an artificial tidal lagoon constructed to drain surrounding settlements and connected to the ocean through a tidal creek and permanent inlet. Little is reported on these mangroves, although the estuary has been the focus of flood mitigation efforts for surrounding neighborhoods (USACE 1987). One informal water quality survey suggested elevated sewage input as evident in high fecal coliform loads and high nutrient concentrations, as well as minimal tidal connectivity and the temporary influence of precipitation on water levels and salinity (USGS 2011). Levittown median annual rainfall from 2012 to 2017 was 1752 mm (this study).

The island’s second largest metropolitan area is Ponce, which falls within the watershed described as lying between the Río Inabón and the Río Loco on the southern Caribbean coast. Unlike the other two watersheds, the mangrove sites at Ponce are largely unconnected and do not share the same estuarine conditions. The three mangrove forests within this study were located at Punta Cabullones, the Rafael Cordero Santiago Port of the Americas, and on the northern shore of Laguna de Salinas at Punta Cucharas. One study has covered the hydrology and water quality of some of these mangroves (Rodríguez-Martínez and Soler-López 2014), which found dissolved oxygen, specific conductance, and salinity to be dependent upon seasonal changes in hydrologic inputs during the wet and dry seasons. The site received a median annual rainfall of 755 mm from 2012 to 2017 (this study).

Based on aerial imagery and previously mentioned hydrological studies, sites have been classified into hydrogeomorphic settings of embayments (i.e. ocean, bay, or lagoon) or canals, and either open or partially restricted to tidal exchange (Figure 1b). Ocean and bay sites by their nature have direct tidal exchange and are thus always classified as open. These sites are the two San Juan Bay sites BAHMIN and BAHMAX, and the ocean site in Ponce, PONMIN. Lagoon or canal sites may have open or partial tidal exchange. Torrecillas lagoon is an open lagoon due to its dredged mouth at boca de cangrejos (Ellis 1976). Open canals are the east end of Suárez canal, SUAMAX, which shares the dredged connection with Torrecillas, the west and dredged portion of the Caño Martin Peña, MPNMIN, and the mouth of the Rio Cocal in Levittown, LEVMIN. In contrast, partially restricted canals are that connecting the Rio Cocal to the Levittown lakes, LEVMID, and the undredged eastern portion of the Caño Martín Peña, MPNMAX. Partially restricted embayments are those of Piñones (PIN) and San José (SANMIN and SANMAX) in the San Juan Bay Estuary, Levittown lakes (LEVMAX) in Levittown, and the Salinas lagoon (PONMID) and the Port of the Americas in Ponce (PONMAX).

### Calculating Urbanization

Spatial datasets used for the selection of study sites and the calculation of urban variables are described in Table 1. All spatial analyses were performed in the R programming language (Yan et al. 2011). Individual functions in R are stated along with their corresponding packages and authors. For brevity, these are reported as package::function, following the notation in the R language. Packages used in these analyses include sp (Bivand et al. 2013), rgeos (Bivand and Rundel 2017), and raster (Hijmans 2016). Commented subroutines for these analyses are provided at github.com/BBranoff/Urban-Mangrove-Hydrology, and a schematic outlining the process is demonstrated in Figure 2.

**Table 1.** Spatial datasets used to determine relative elevations and to quantify the urbanization surrounding each study site. Variables were sampled from a sampling area described by a circle of radius 500 m and centered on individual study sites.

| Variable | Dataset Description | Source |
| --- | --- | --- |
| **Land Cover**<br>**Urban Vegetation and Water** | 2 m resolution land cover raster for Puerto Rico in 2010 | (Office for Coastal Management 2017) |
| **Mangrove Coverage** | 30 m resolution continuous mangrove coverage raster for 2012 | (Hamilton and Casey 2016) |
| **Population Density** | Total population shapefile for 2010 in Puerto Rico by census block | (U.S. Census Bureau 2010) |
| **Road Length** | Road network shapefile for Puerto Rico in 2015 | (U.S. Census Bureau 2015) |
| **Elevation** | Digital elevation models for sites, at 3 & 1 m horizontal resolution and 9 and 4.2 cm vertical accuracy for the northern and southern sites, respectively | (NOAA 2015a, b) |

**Figure 2.**
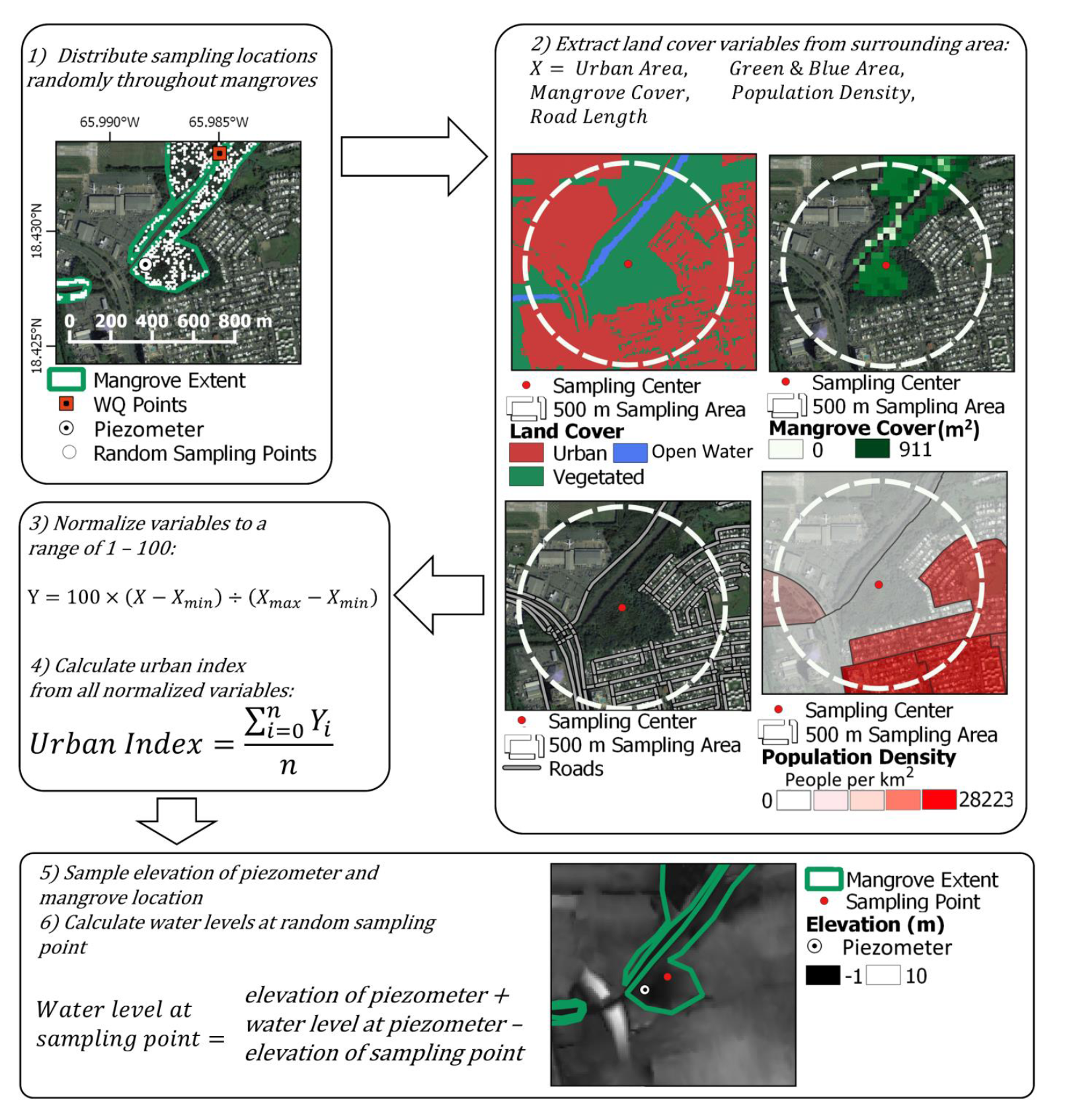
A schematic of the spatial sampling routine for the calculation of the urban index and water levels at random mangrove locations throughout the study area. Sampling locations were distributed randomly throughout the mangrove habitat. Land cover within the surrounding 500 meters of these locations was then sampled and used to calculate the urban index. Elevations were sampled at each location and at the nearest piezometer, which were used to calculate the water levels at all locations.

To begin, one hundred random points were placed within mangrove classified habitat in each watershed of Puerto Rico (NOAA Biogeography Program 2011). To assess the surrounding land coverage, a sampling circle of radius 500 m was created around each point, and mangrove coverage and land cover datasets were then cropped and masked to include only those cells within the circle. The total mangrove coverage for each point was then calculated by summing the mangrove cell values. Land cover was calculated by first counting the number of cells within each class, in which urban was classes 2 and 5 (Impervious and Developed Open Space, 1 respectively), and green and blue area was the sum of all vegetation and open water classes, defined as classes 6 through 18, and 21. These sums were then multiplied by the area of each cell, which had a mean of 2.01 m.

Road lengths within each circle were calculated by first clipping the entire road network to include only the area within the circle and summing the combined length. Population density within the sampling circle was calculated using 2010 US Census data at the block level. The total population within the sampling circle was calculated assuming all people lived in non-road impervious surfaces. The area of non-road impervious surfaces within each block was calculated by first obtaining the total area of impervious surfaces through the methods of the previous paragraph, using the boundary of each census block to sample the land cover within. This was then used to calculate the population density per unit area of non-road impervious surfaces for each block, which was then multiplied by the area of non-road impervious surfaces within each part of each census block contained in the circular sampling area. This resulted in the total number of people within the sampling area, which was then divided by the area of the circle to give the population density.

These variables were used independently in further analyses, along with a combination of all of them representing an urban index at each location. The index was calculated based upon a similar method for aquatic ecosystems (McMahon and Cuffney 2000). The urban index value is calculated using the following equation:

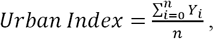

where *n* is the number of variables used in the index, in this case the five variables of urban area, green and blue area, mangrove area, populations density, and road length. *Y*_*i*_ represents the variables normalized to a range of 0 to 100 through the following equation:

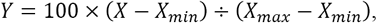

In which *Y* and *X* represent the normalized and raw values, respectively. In the case of mangrove and blue-green coverage, the normalized variable was reversed by subtracting from 100, so that lower values represent more urbanness. The urban index is thus a representation of the relative intensity of urbanization, in which 100 is the most urban site and 1 is the least urban site.

### Water Level and Weather Data Acquisition

Water levels models were constructed from water level recordings made throughout the study area and from precipitation and barometric pressure observations from nearby weather stations. The models covered a five-year period, from June 1, 2012 to June 1, 2017. Data acquisition and subsequent modeling and statistical analyses were done in the R programming language (Yan et al. 2011). Individual functions with R are stated along with their corresponding packages and authors. For brevity, these are sometimes reported as package::function, following the notation in the R language. Additional utilized packages not mentioned below are the zoo (Zeilis and Grothendieck 2005), RJSONIO (Temple Lang 2014), and HyetosMinute packages (Kossieris et al. 2016).

#### Water Levels

Water levels were recorded in 16 locations within the three watersheds in Puerto Rico (Figure 1b). Piezometer wells were constructed from 2 m segments of 3” diameter PVC tubing. One half of the length of the well was perforated with 1 mm x 1 cm slits placed every 1 cm using a sanding disc attached to a Dremel rotary tool. The wells were caped with PVC caps and one Onset HOBO U20l water level logger was hung from a 1.75 m cable attached to the top cap. Wells were placed in holes excavated at the shoreline using a 3” corer to a depth of 1.5 m. The water level loggers were programmed to take pressure and temperature readings every 15 minutes. Data were extracted using an Onset HOBO waterproof shuttle at varying intervals averaging once every three months. Water levels were calculated from pressure readings using the HOBO barometric compensation tool in the HOBO Pro software and atmospheric pressure readings recorded at varying locations as described below.

#### Weather Data

Site precipitation and atmospheric pressure were acquired from varying sources as detailed in Table 2 and assembled into one-hour observations over the five-year period. Hourly observations were collected from wunderground.com using their Weather API feature (Table 2). For San Juan (SJU), this represented a complete set of observations, but the two other locations required additional data sources and approximations to compensate for large gaps in the wunderground data. Missing precipitation data were filled using daily values from the National Climate Data Center at various stations (Table 2). Hourly rainfall from these stations was reconstructed from the daily data through the *DisagSimul* function. Missing barometric pressure observations were approximated using a linear interpolation between known values calculated through the *na.approx* function.

**Table 2.**
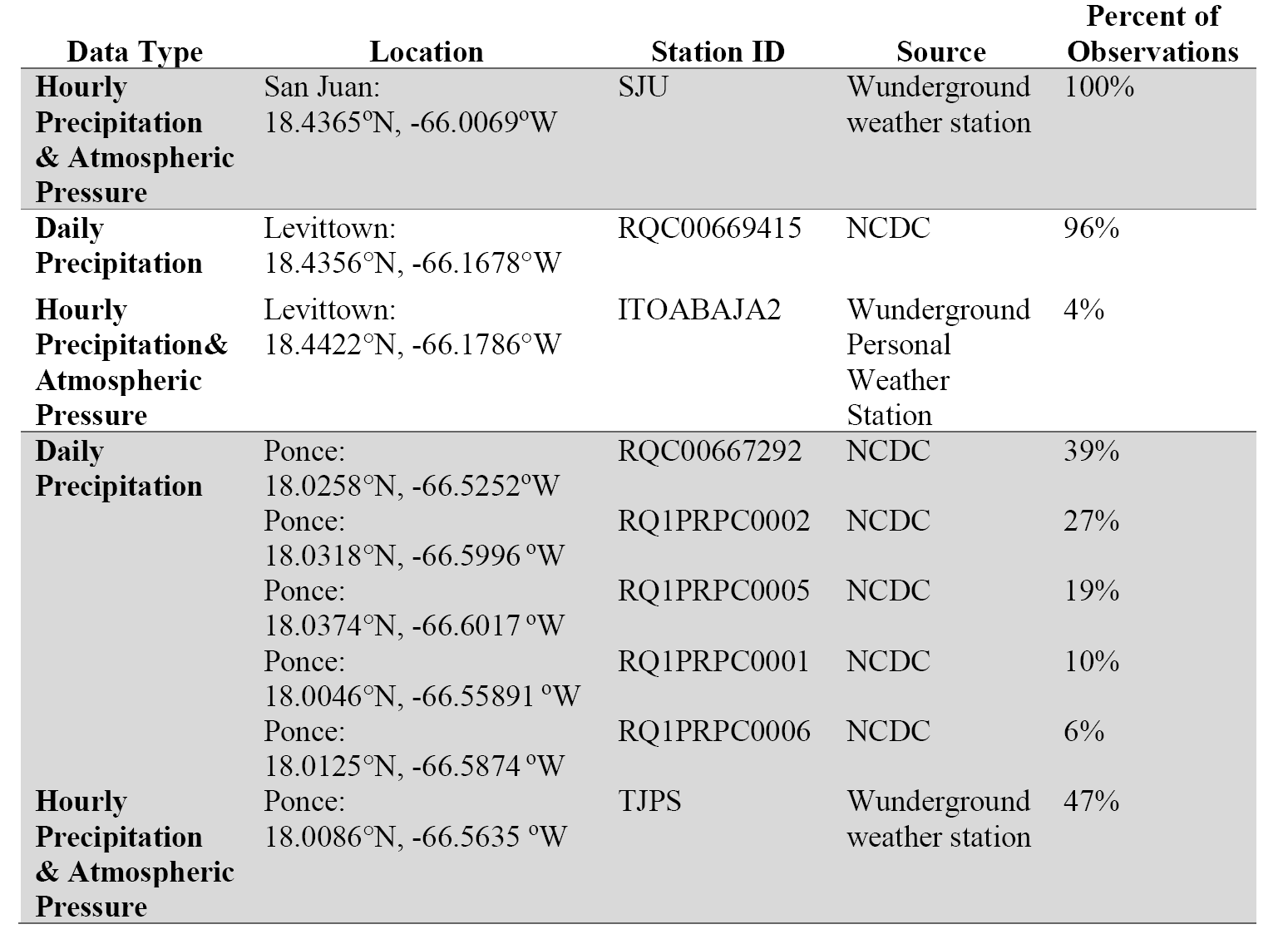
Weather data sources for the three regions included in the study (San Juan, Ponce, and Levittown, Puerto Rico). NCDC is the National Climate Data Center. Percent of observations are the percent of total rain observations for each watershed that were obtained from the corresponding sources.

#### Surface Water Chemical Properties

Surface water chemical properties were only available for the San Juan Bay Estuary, where most of the study sites are found. Measurements were downloaded from the San Juan Bay Estuary Program’s water quality monitoring program. Monthly metrics were pH, temperature (^°^C), dissolved oxygen (DO) (mg/L), Salinity (PSS), and turbidity (NTU). Biannual metrics were total Kjeldahl nitrogen (mg/L), total nitrate and nitrite (mg/L), total phosphorous (mg/L), total organic carbon (TOC) (mg/L), ammonia (mg/L), biological oxygen demand (BOD) (mg/L), and oil and grease (mg./L). The lowest detection limits and total number of samples for each metric at each study site are included as Table 3. Although mangrove nutrition would be more accurately represented by pore-water chemistry, there is a documented relationship between surface and pore-water in mangrove systems (Bouillon et al. 2007; Gleeson et al. 2013), and also between surface water chemistry and mangrove distribution (Sherman et al. 1998). Thus, these relatively easier to obtain surface water measurements will serve as a good indication of the mangrove chemical environment over a large area.

**Table 3.**
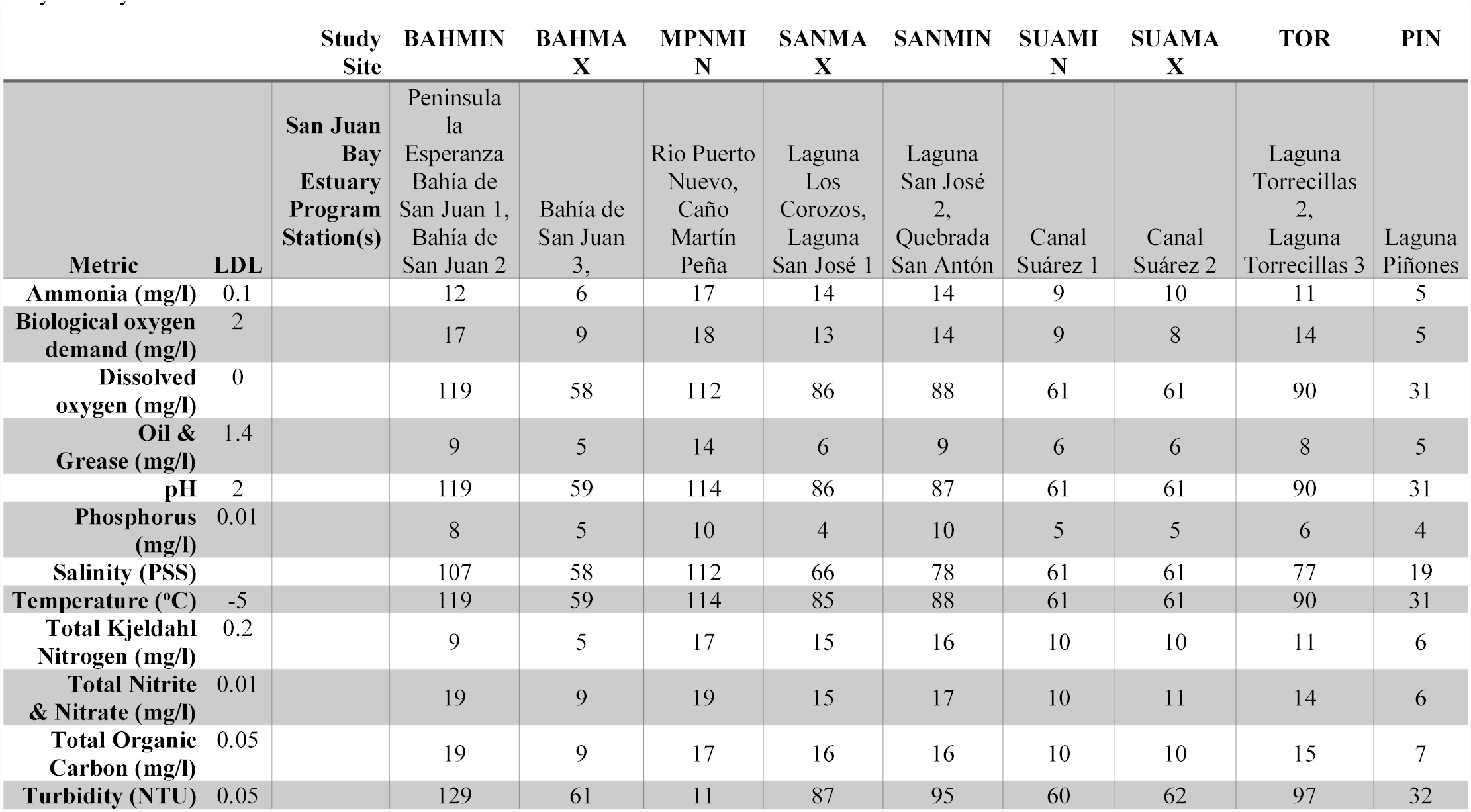
Lowest detection limits (LDL) and sample sizes for surface water chemistry measurements taken from the San Juan Bay Estuary Program corresponding to the study period: June 1, 2012 to June 1, 2017. There is no station for the MPNMAX study site or for those outside of the San Juan Bay Estuary.

### Modeling

As with all other analyses, water level modeling was done in the R programming language and a detailed, commented copy of the code can be found at github.com/BBranoff/Urban-Mangrove-Hydrology. In addition to the packages mentioned below, the TTR (Ulrich 2017), zoo (Zeilis and Grothendieck 2005), minqa (Bates et al. 2014), and relaimpo packages (Grömping 2006) were also used throughout the process. Water levels at each location were modeled using a combination of tidal and precipitation inputs. Tidal components were modeled through the *ftide* function in the TideHarmonics package (Stephenson 2016), which uses timestamped water level observations to compute model coefficients for up to 409 harmonic tidal constituents. Constituent amplitudes and phases interact to give a change in water level, relative to the mean. Water levels due to tides at a given time, *Tides*(*t*), can thus be modeled from all *N* harmonic constituents by summing each individual *n* constituent using the following equation from Stephenson (2016):

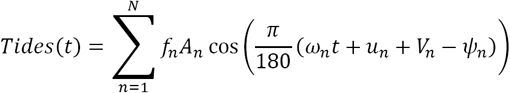

The amplitude, angular frequency, equilibrium phase, phase lag, and nodal corrections for each constituent are given by *A*_n_, *ω*_n_, *V*_n_, *Ψ*_n_, *u*_n_ and *f*_n_ respectively. The time (*t*) is represented as the length, in minutes, from the origin of the model, which is dependent upon the period of observations at each site. Models were calculated for all observations in which the four-day cumulative rainfall was less than 1 mm to avoid interference from precipitation inputs. The resulting coefficients of each tidal constituent were saved for further analyses and are given in the Appendix A. These models were then used with the *predict.tide* function to predict water levels every hour over the five-year period. The result is water level predictions based only on tidal components.

Tidal predictions were then used along with rainfall data and the observed water levels to compute contributions from precipitation. This contribution was modeled at each location through a combination of moving averages and moving sums, as has been implemented in other studies (Dawson and Wilby 1998; Toth et al. 2000; Altunkaynak 2007). Short-term response to rainfall was modeled with an exponential moving average (EMA) of precipitation. The short-term contribution from precipitation is thus given as:

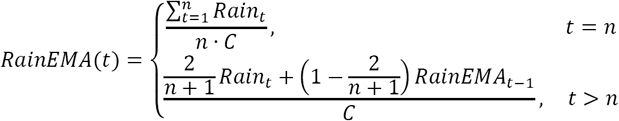

In which *Rain*_t_ is the observed precipitation at time t, *n* is the moving average window, and *n* is a dampening constant. Long-term precipitation input was modeled using a moving sum of rainfall data in the form:

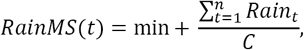

In which min is the baseline water level, taken to be the minimum recorded observation, *Rain* is the observed precipitation in mm, *n* is the moving sum window, and *C* is a dampening constant. Coefficients for both short and long term rainfall contributions were optimized for each location using a BOBYQA optimization, by minimizing the residual sum of squares between the observed water levels and the sum of the tidal and rain models (Powell 2009).

Using the above equations, the water level at each location at any given time can be predicted by combining the tidal and precipitation components through the equation:

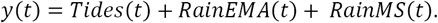

In some cases, optimization resulted in zero contribution from tidal, or short and long-term rainfall contributions. All tidal and rainfall model coefficients are provided in Appendix B.

With both the tidal and precipitation contributions to water levels modeled for each piezometer, their relative importance of each in the overall model was also calculated. This was done for individual tidal constituents and for the combined rainfall constituent. Relative importance was calculated as the contribution of each constituent to the R^2^ of the full model against the observations (Lindeman 1980). This importance value was calculated for the ten tidal constituents with the greatest amplitude. The individual and cumulative R^2^ of all constituents in the water level models are reported in the results.

### Mangrove Elevations

The above water level models were used to predict water levels throughout the mangroves in the three watersheds of the study. This was accomplished by assuming a planar, not curved water surface from tidal asymmetry (Wolanski et al. 1993). Thus, water levels at random points were calculated by subtracting their elevation from that of the observation piezometers and applying this adjustment to the predicted water levels (Figure 2). Mangroves were defined by a benthic habitat map of Puerto Rico (NOAA Biogeography Program 2011), and elevation was taken from two 3 m horizontal resolution coastal digital elevation models (DEM), one for San Juan and Levittown (NOAA 2015a), and another for Ponce (NOAA 2015b). As with other analyses, a copy of the R subroutines is provided on GitHub at github.com/BBranoff/Urban-Mangrove-Hydrology. Additional packages utilized but not mentioned below are deldir (Turner 2017), sp (Bivand et al. 2013), and raster (Hijmans 2016).

One thousand random points were generated within each of the sixteen mangrove zones (Figure 1b), and their elevations extracted from the DEMs. There were occasional small discrepancies between the benthic habitat map and the DEMs, resulting in extreme and likely mistaken elevations assigned to mangrove habitat. To remove these extreme values, the randomly sampled points were limited to elevations between the 10% and 90% quantiles. The resulting elevations were subtracted from that at the piezometer location, and this adjustment was added to the predicted water levels to give a time series of water levels at each point at every hour within the five-year period.

### Calibration and Validation

Models were calibrated using the above described equations and optimization techniques on the first 90% of observations, withholding the last 10% for validation. The optimized models were then validated by predicting water levels for the remaining 10% and comparing to the observations. This was done through visual inspection and by calculating the R^2^ between the two. Additionally, the mean absolute error (ε) between observations and predictions was calculated as,

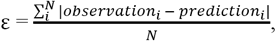

for all *N* validation observations at each site. An error of less than 0.1 m is recommended for estuarine waters (Williams and Esteves 2017), and a combination of both visual and statistical validation is recommended for gauging model accuracy (Biondi et al. 2012). Because the models presented here are not intended to be used for engineering or design input, but rather for ecological studies, a satisfaction criterion of R^2^ > 0.5 and ε < 0.1 m was implemented. Hydrographs of observed and predicted water levels were also inspected to ensure satisfactory agreement. Models not meeting these criteria in both calibration and validation periods are thus reported on but excluded from hypothesis testing.

### Flooding Parameters

The resulting time series of water levels were analyzed for daily flooding parameters of mean flood duration, mean dry duration, mean flood frequency, average depth, and percent of time flooded. The same parameters were calculated for each month in the time series to correspond to monthly water quality measurements. Water levels were separated into flood (≥ 0) and dry (< 0) conditions, and analyzed for length, frequency, and mean at the daily and monthly level.

### Statistical Tests

Differences in predictor variables of flooding dynamics and water quality metrics between mangrove zones were identified through ANOVA using the *aov* function, and subsequent post-hoc differences through a Tukey honest significance test by the *TukeyHSD* function, both from base R (Yan et al. 2011). In testing mean differences between urban and non-urban sites, defined as those with and without any urban area within the 0.5 km sampling radius, student t-tests were used through the *t.test* function of base R. Linear models were constructed through the *lm* function in base R using the form y∼x and y∼log(x). Results were plotted using the *ggplot* function from the ggplot2 package and linear and logarithmic models through the *stat_smooth* function (Wickham 2009).

## RESULTS

### Model Accuracy and Constituent Importance

Water level model fits varied across the systems (Figures 3 & 4). For the calibration period, all models explain fifty percent or more of the variation in observed water levels (R^2^) and eight (57%) of the models explain seventy percent or more of the variation in water levels (Figure 3b & 4b). Mean absolute error (ε) among all models during the calibration period was 7.4 cm (Figure 3b & 4b). Validations for all models performed marginally worse than calibrations, with only twelve of the models explaining fifty percent or more of the variation in water levels, and a mean absolute error of 4.4 cm (Figure 3c & 4c). Two models, LEVMAX and PONMAX, did not meet the criteria set for satisfactory agreement with observations in both calibration and validation periods (R^2^ > 0.5, ε < 0.1). These sites are thus reported on in figures as faded symbols and are excluded from further hypotheses testing involving flooding dynamics.

**Figure 3.**
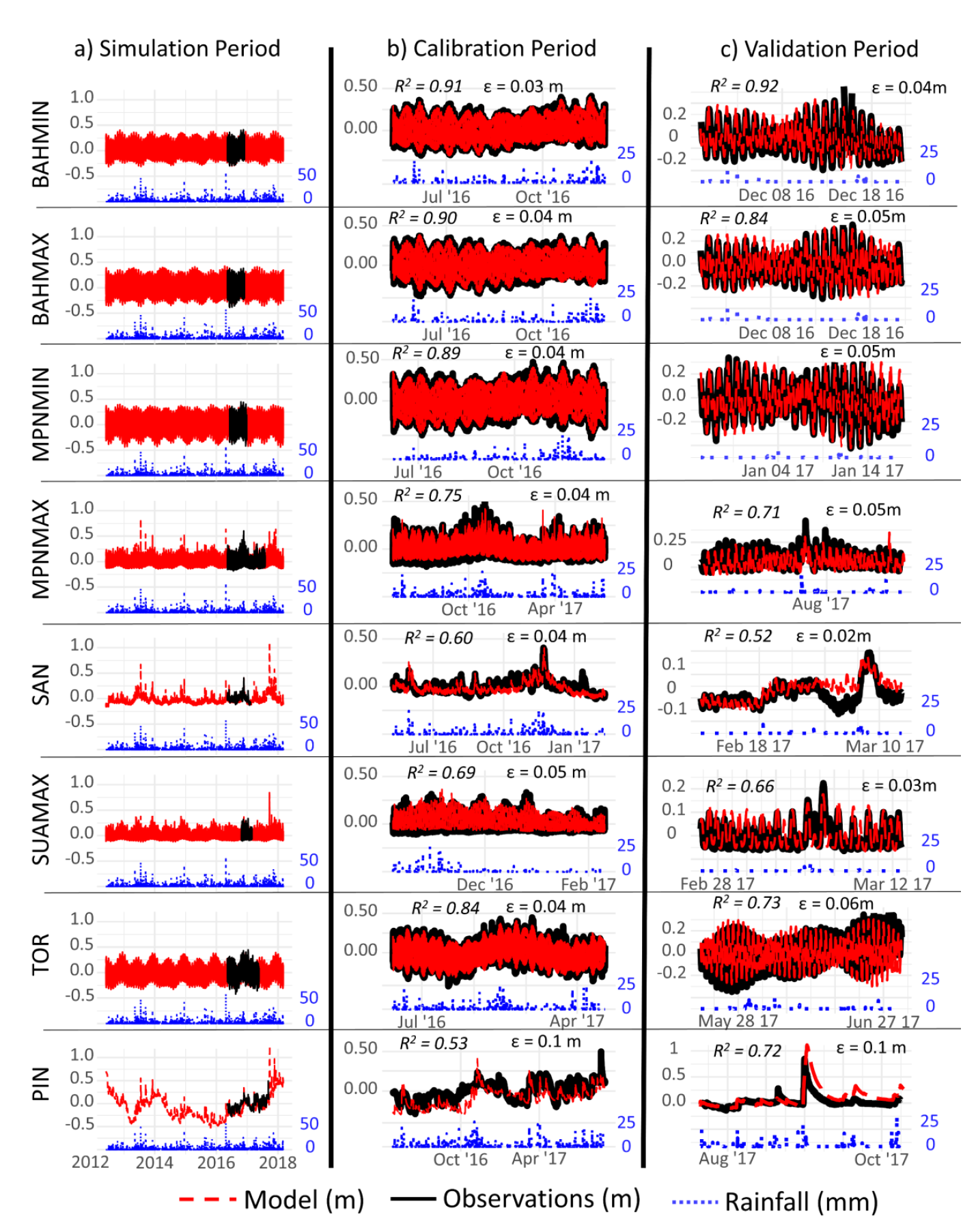
Rainfall, water level observations, and water level models in the San Juan Bay Estuary for the entire length of the analysis, 2012 – 2017 (a), for the observation period at each location (b), and for the validation period (c). Water levels are in reference to the mean and are plotted on the right axis. Rainfall is hourly and is plotted on the right axis. Water levels in (a) and (b) are on a fixed scale to highlight the difference in ranges between locations. Water level ranges in all other panels are unique for each location. Observations and models for SANMIN, SANMAX and SUAMIN were nearly identical, and are thus here referred to as SAN.

Constituent influences on the water levels varied in importance across the studied systems (Table 4, Table 5). The following percentages represent the contributions of each constituent, or group of constituents, to the observed variations in water levels. In the northern sites of the San Juan Bay Estuary and in Levittown, the three most important constituents were the principal lunar semi-diurnal (M2), two lunar diurnals (K1 & O1). Fifty-three, and fifty-four percent of the water level variations in Piñones (PIN) and San José Lagoons (SAN), respectively, were explained by the short and long-term precipitation models and the water levels at PIN could not be modeled using tidal constituents. In contrast, in the San Juan Bay (BAHMIN & BAHMAX) and in the dredged portion of the Caño Martin Peña (MPNMIN), precipitation explained less than 1% of the variation in water levels. In Levittown, the K1, M2 and O1 constituents combined and along with rainfall explained 45% and 54% of the variation in water levels at LEVMIN and LEVMID, respectively. Most of the remaining variation in water levels at these two sites is explained by the luni-solar synodic fortnight (Msf) and the luni-monthly (Mm). Fifty-eight percent of the variations in the LEVMAX station were explained only by precipitation and no tidal model could be constructed. At Ponce, the three most important constituents were the two solar diurnals (S1 & P1) and rain. These constituents explained 27%, 57%, and 31% of the water level variations at PONMIN, PONMID, and PONMAX, respectively. Rain explained 56% and 31% of the water level variations at PONMID and PONMAX, respectively, and less than 1% at PONMIN.

**Table 4.** Water level model constituents and their importance in explaining variations in the water levels of the selected waterbodies of the San Juan Bay Estuary. The overall rank is that of all waterbodies combined, and the individual ranks are those for each individual water body. “% Exp.” is the mean contribution to the explained variance of the observed water levels in iterative models with varying constituent orders. “Cum. % Exp.” is the cumulative percent of the explained variation in the observed water levels for a model containing the constituent and all above constituents. SAN represents sites of SANMIN, SANMAX and SUAMIN as all three models were nearly identical. For the San Juan Bay estuary, rain is the fifth most important constituent in explaining water level variations, but overall the principal lunar semi-diurnal tidal constituent (M2) is most important.

| CONSTITUENT | Overall Rank | BAHMAX |  |  | BAHMIN |  |  | MPNMIN |  |  | MPNMAX |  |  |
| --- | --- | --- | --- | --- | --- | --- | --- | --- | --- | --- | --- | --- | --- |
|  |  | Rank | % Exp. | Cum. % Exp. | Rank | % Exp. | Cum. % Exp. | Rank | % Exp. | Cum. % Exp. | Rank | % Exp. | Cum. % Exp. |
| M2 | 1 | 1 | 59% | 59% | 1 | 55% | 55% | 1 | 57% | 57% | 1 | 28% | 28% |
| K1 | 2 | 2 | 15% | 74% | 2 | 14% | 70% | 2 | 13% | 70% | 2 | 12% | 40% |
| O1 | 3 | 3 | 9% | 83% | 3 | 9% | 79% | 3 | 9% | 79% | 4 | 9% | 49% |
| RAIN | 4 | 10 | 0% | 83% | 10 | 0% | 79% | 10 | 0% | 79% | 3 | 10% | 60% |
| N2 | 5 | 4 | 2% | 85% | 5 | 2% | 81% | 5 | 3% | 82% | 7 | 2% | 61% |
| P1 | 6 | 11 | 0% | 85% | 11 | 0% | 81% | 6 | 2% | 84% | 8 | 1% | 63% |
| S2 | 7 | 7 | 1% | 86% | 7 | 1% | 82% | 7 | 1% | 85% | 10 | 1% | 63% |
| Mf | 8 | 12 | 0% | 86% | 9 | 0% | 82% | 9 | 0% | 85% | 15 | 0% | 63% |
| Sa | 9 | 19 | 0% | 86% | 6 | 2% | 85% | 16 | 0% | 85% | 5 | 6% | 69% |
| Q1 | 10 | 14 | 0% | 86% | 15 | 0% | 85% | 12 | 0% | 85% | 13 | 0% | 69% |
| ALL | - | - | - | 91% | - | - | 90% | - | - | 89% | - | - | 75% |
|  |  | SAN |  |  | SUAMAX |  |  | TOR |  |  | PIN |  |  |
| M2 | 1 | 2 | 1% | 1% | 1 | 34% | 34% | 1 | 46% | 46% | - | - | - |
| K1 | 2 | 3 | 1% | 2% | 2 | 11% | 44% | 2 | 11% | 58% | - | - | - |
| O1 | 3 | 4 | 1% | 2% | 4 | 7% | 51% | 4 | 7% | 65% | - | - | - |
| RAIN | 4 | 1 | 57% | 59% | 3 | 10% | 61% | 8 | 1% | 66% | - | - | - |
| N2 | 5 | 7 | 0% | 59% | 9 | 1% | 63% | 6 | 2% | 68% | 1 | 1 | 53% |
| P1 | 6 | 6 | 0% | 60% | 7 | 2% | 64% | 7 | 2% | 70% | - | - | - |
| S2 | 7 | 15 | 0% | 60% | 10 | 1% | 65% | 9 | 1% | 71% | - | - | - |
| Mf | 8 | 5 | 0% | 60% | 22 | 0% | 65% | 10 | 0% | 71% | - | - | - |
| Sa | 9 | 19 | 0% | 60% | 8 | 2% | 67% | 5 | 5% | 76% | - | - | - |
| Q1 | 10 | 8 | 0% | 60% | 19 | 0% | 67% | 11 | 0% | 76% | - | - | - |
| ALL | - | - | - | 60% | - | - | 72% | - | - | 84% | - | - | 53% |

K1 is the lunar diurnal. M2 is the principal lunar semidiurnal. Mf is the lunar fortnightly. N2 is the larger lunar elliptic semidiurnal. O1 is the lunar diurnal. P1 is the principal solar diurnal. Q1 is the larger lunar diurnal elliptical. S2 is the solar semidiurnal. Sa is the solar annual.

**Table 5.** Water level model constituents and their importance in explaining variations in the water levels of the selected waterbodies of Levittown and Ponce. The overall rank is that of all waterbodies combined, and the individual ranks are those for each individual water body. “% Exp.” is the mean contribution to the explained variance of the observed water levels in iterative models with varying constituent orders. “Cum. % Exp.” Is the cumulative percent of the explained variation in the observed water levels for a model containing the constituent and all above constituents. In levittown, rain is the tenth most important constituent in explaining water level variations, and overall the principal lunar semi-diurnal tidal constituent (M2) is most important. In Ponce, Rain is the second most important constituent, behind the principal solar semidiurnal (S1).

| CONSTITUENT | LEVMIN |  |  |  | LEV MID |  |  | LEVMAX |  |  |
| --- | --- | --- | --- | --- | --- | --- | --- | --- | --- | --- |
|  | Overall Rank | Rank | % Exp. | Cum. % Exp. | Rank | % Exp. | Cum. % Exp. | Rank | % Exp. | Cum. % Exp. |
| **K1** | 1 | 1 | 18% | 18% | 2 | 17% | 17% | - | - | - |
| **M2** | 2 | 4 | 11% | 29% | 1 | 24% | 41% | - | - | - |
| **O1** | 3 | 3 | 11% | 40% | 3 | 13% | 54% | - | - | - |
| **1M.1P1** | 4 | - | 0% | 40% | 4 | 5% | 59% | - | - | - |
| **MSF** | 5 | 2 | 13% | 53% | 11 | 0% | 59% | - | - | - |
| **RHO1** | 6 | 7 | 3% | 55% | 7 | 0% | 59% | - | - | - |
| **S2** | 7 | 8 | 2% | 58% | 6 | 2% | 61% | - | - | - |
| **RAIN** | 8 | 6 | 5% | 63% | 9 | 0% | 61% | 1 | 58% | 58% |
| **MF** | 9 | - | 0% | 63% | 8 | 0% | 61% | - | - | - |
| **MM** | 10 | 5 | 9% | 72% | 14 | 0% | 61% | - | - | - |
| **ALL** |  | - | - | 75% | - | - | 64% | - | - | 58% |
| CONSTITUENT | PONMIN |  |  |  | PONMID |  |  | PONMAX |  |  |
|  | Overall Rank | Rank | % Exp. | Cum. % Exp. | Rank | % Exp. | Cum. % Exp. | Rank | % Exp. | Cum. % Exp. |
| **P1** | 1 | 4 | 5% | 5% | 3 | 1% | 1% | - | 0% | 0% |
| **RAIN** | 2 | 10 | 0% | 5% | 1 | 56% | 57% | 1 | 31% | 31% |
| **S1** | 3 | 2 | 22% | 27% | 7 | 0% | 57% | - | 0% | 31% |
| **K1** | 4 | 1 | 28% | 55% | 4 | 0% | 57% | 10 | 0% | 31% |
| **O1** | 5 | 3 | 19% | 74% | - | 0% | 57% | 8 | 0% | 31% |
| **SSA** | 6 | 13 | 0% | 74% | 2 | 19% | 77% | 3 | 6% | 37% |
| **MSF** | 7 | 9 | 0% | 74% | - | 0% | 77% | 4 | 0% | 37% |
| **SA** | 8 | 6 | 2% | 76% | 12 | 0% | 77% | 2 | 30% | 67% |
| **Q1** | 9 | 5 | 2% | 78% | 9 | 0% | 77% | - | 0% | 67% |
| **S2** | 10 | 4 | 0% | 78% | 6 | 0% | 77% | 6 | 0% | 67% |
| **ALL** |  | - | - | 80% | - | - | 77% | - | - | 67% |

K1 is the lunar diurnal. M2 is the principal lunar semidiurnal. MF is the lunisolar fortnightly. MM is the lunar monthly. MSF is the lunisolar synodic fortnightly. N2 is the larger lunar elliptic semidiurnal. O1 is the lunar diurnal. P1 is the solar diurnal. RHO1 is a diurnal. S1 is the solar diurnal. S2 is the principal solar semidiurnal. SA is the solar annual. SSA is the solar semi-annual. PSI1 is a diurnal.

### Flooding Dynamics

Random mangrove locations throughout the study sites varied in their flooding dynamics (Figure 5). Median average depths in San Juan, Levittown, and Ponce were -1.4, -6.0, and 0.08 cm, respectively, and means were significantly different among all watersheds (ANOVA; p < 0.05). Ponce depths were 8.9 cm greater than those at Levittown (ANOVA; p < 0.001), and 5.8 cm greater than those at San Juan (p < 0.001). San Juan depths were 3 cm higher than Levittown (ANOVA; p < 0.05). Median flood length in the three systems were 0.64, 0.17, and 2.3 days, respectively. San Juan mean flood length was 108 days longer than Levittown and 102 days longer than Ponce (ANOVA; p<0.001). Median daily flood frequency, in which 0 is either constantly flooded or constantly dry, were 1.0, 0.7, and 1.1 respectively in San Juan, Levittown and Ponce. On average San Juan mangroves flooded 0.3 and 0.2 more times per day than in Levittown and Ponce (ANOVA; p < 0.001), corresponding to one extra flood every 3 and 5 days, respectively. Mangroves of Ponce flooded 0.2 times per day more than those of Levittown, corresponding to one extra flood every 5 days (ANOVA; p < 0.01). Median percent of time flooded was 40,11, and 48%, respectively, in San Juan, Levittown and Ponce. San Juan was flooded 13% more of the time that Levittown was flooded and 5.8% less than Ponce (p < 0.001). Ponce was flooded 19% more of the time than Levittown (p < 0.001).

**Figure 4.**
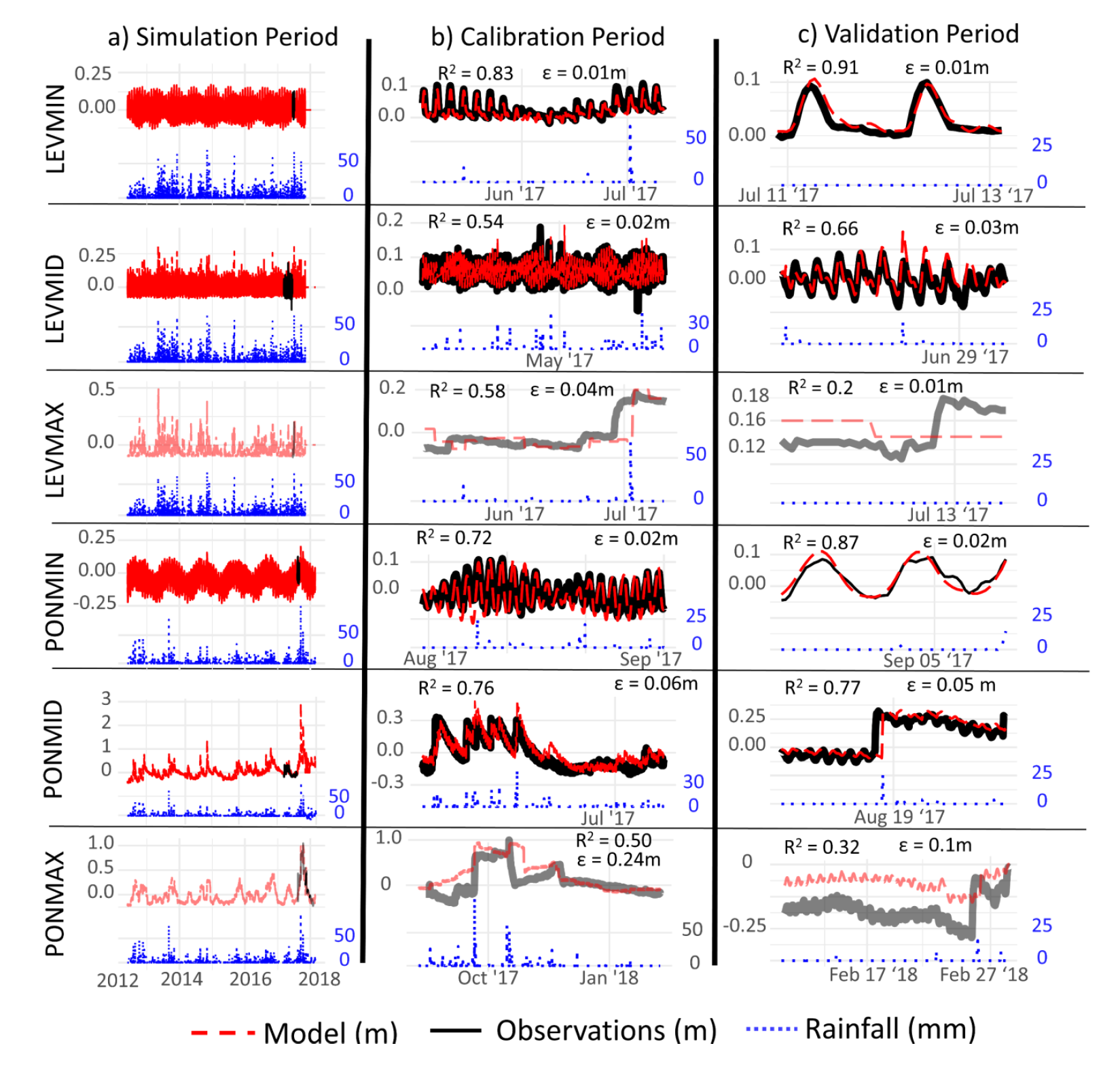
Rainfall, water level observations, and water level models in Levittown and Ponce for the entire length of the analysis, 2012 – 2017 (a), for the observation period at each location (b), and for the validation period (c). Water levels are in reference to the mean and are plotted on the right axis. Rainfall is hourly and is plotted on the right axis. Water levels in (a) and (b) are on a fixed scale to highlight the difference in ranges between locations. Water level ranges in all other panels are unique for each location. Models for LEVMAX and PONMAX did not meet validation criteria and are thus excluded from future hypothesis testing.

**Figure 5.**
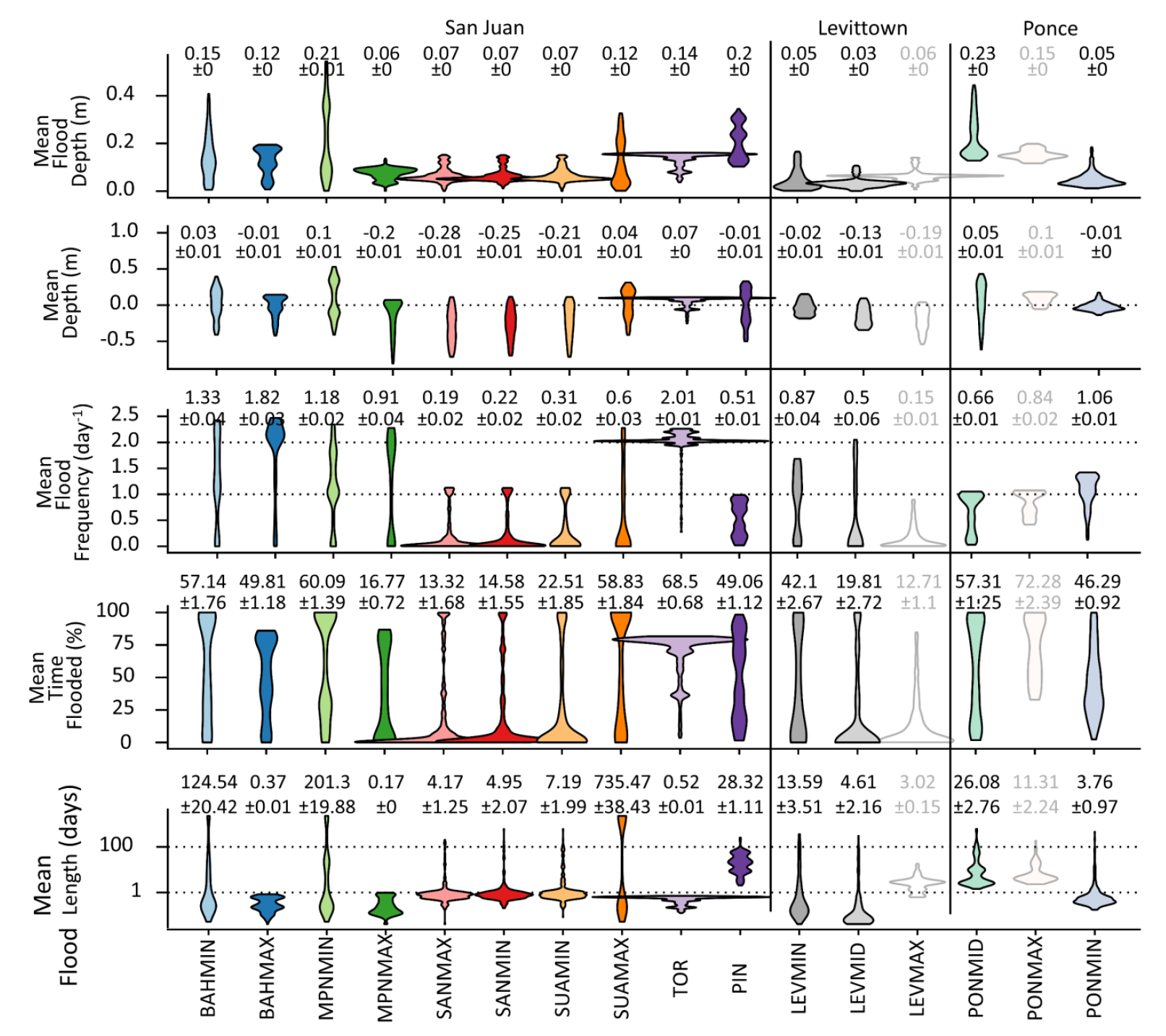
Violin plots of mean flooding metrics from random locations within the mangrove zones over the course of the five-year modeling period. Annotations are means and standard errors for each location. Locations are ordered from west to east in each watershed. Constantly flooded or dry locations have a flood frequency of 0. Shapes of the violin plot are determined by value densities, with the widest portion of the “violin” being representative of the highest density of values. PONMAX, PIN, LEVMAX, and PONMID are most distinct and characterized by long hydroperiods with minimal daily flood frequencies.

Among sites, flooding metrics were variable, and no single site was distinct in all metrics. The dredged portion of Caño Martin Peña (MPNMIN) held the greatest average depth at 10 cm, which was significantly different than all other sites except Torrecillas lagoon (TORMIN) (ANOVA; mean difference = 17 cm, p<0.001). In flooded depth, PONMID was greatest at 22.5 cm, which was significantly greater than all other sites except MPNMIN (ANOVA; mean difference = 12 cm, p <0.001). In contrast, SANMIN experienced the lowest average depth at - 24.6 cm (24.6 cm above water), which was significantly different than all sites except MPNMAX, SANMIN and SUAMIN (ANOVA; mean difference = 21 cm, p < 0.001).

Piñones lagoon was exceptional in hydroperiod (flood length), with a median of 20.5 days, five times longer than PONMID and at least twenty times longer than the other sites. However, SUAMAX held the greatest mean (not median) flood length at 735 days, longer than all other sites (ANOVA; mean difference = 703 days, p <0.001). In contrast, MPNMAX exhibited the shortest hydroperiod of 0.2 days, which was lower than all other sites except LEVMID (ANOVA; mean difference = 8 days, p = <0.001). Flood frequency was lowest at SANMAX at 0.2 day, which was lower than all other sites except SANMIN and SUAMIN (ANOVA; mean difference = 0.7 per day, p < 0.001). The highest flood frequency was shown by TORMIN at 2 per day, higher than all other sites (ANOVA; mean difference = 1.2 per day, p < 0.001). In average depth, MPNMIN was the greatest at 0.1 m and greater than all other sites (ANOVA; mean difference = 0.17 m, p < 0.001), and SANMAX was the least at -0.28 m (0.28 m above water), lower than all other sites except SANMIN (ANOVA; mean difference = 0.24, p < 0.001).

### Surface Water Chemical Properties

In Tukey honest significant difference tests, Piñones (PIN) was singled out in surface water chemical properties (Figure 6). This lagoon was characterized by significantly greater total Kjeldahl nitrogen (mean difference = 8.6 mg/L; p < 0.001), TOC (mean difference = 12.9 mg/L; p < 0.001), turbidity (mean difference = 176.2 NTU; p < 0.001), and BOD (mean difference = 15.5 mg/L; p < 0.01) than all other water bodies, as well as significantly higher temperature than half of the other stations (mean difference = 1.7 ^°^C; p < 0.05). The BAHMAX station was also distinct in having higher salinity than all other stations (mean difference = 16.7 PSU; p < 0.001). SANMIN held the highest mean nitrate and nitrite concentration but was only significantly different than BAHMIN (difference = 0.3, p < 0.05) and SANMAX (difference = 0.3, p < 0.05). SANMAX held the highest pH, which was significantly different than SUAMAX (difference = 0.3, p < 0.01), TOR (difference = 0.3, p < 0.01), and PIN (difference = 0.4, p < 0.05). There were no differences in total phosphorus or oil and grease concentrations between sites.

**Figure 6.**
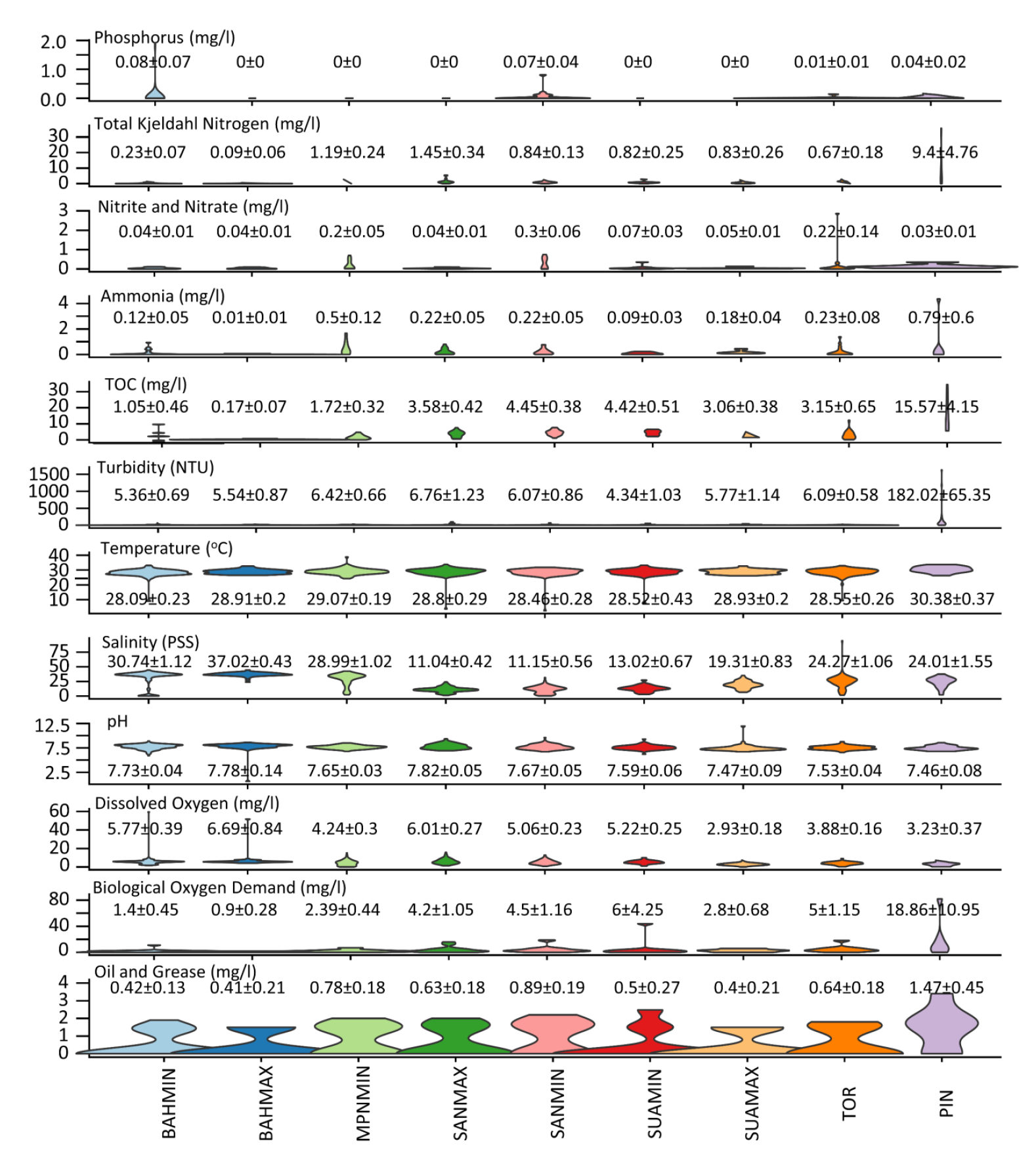
Violin plots of water chemistry throughout the San Juan Bay Estuary. Annotations are means and standard errors for each location. Nitrogen concentrations were abnormally high in some of the waterbodies, which may be associated with sewage effluent. PIN was most distinct, with significantly higher TOC, Kjeldahl nitrogen, and temperature in comparison to the other water bodies.

### Land cover, Surface Water Chemistry and Flood Dynamics

Surface water chemistry metrics resulted in numerous significant models when predicted by surrounding land cover (Figure 7). All metrics except nitrate and nitrite, total phosphorus (P), dissolved oxygen (DO), pH and salinity resulted in significant models when predicted by surrounding land cover. However, almost all models were significant only because of an outlier site, PIN (Piñones lagoon), and could not be repeated when this site was removed from the analysis. Biological oxygen demand was the only metric to be significantly modeled with or without the PIN site, and resulted in a negative relationship with surrounding road length (R^2^ = 0.59, p < 0.05). Otherwise, trends suggest decreasing nitrogen, carbon, turbidity, oil, and temperature with increasing urbanization. Salinity was found to significantly increase with flood frequency (linear model; R^2^ = 0.55, p < 0.05), and oil and grease concentrations were found to increase with flooding duration and proportion of time flooded (linear models; R^2^ = 0.5, p < 0.05).

**Figure 7.**
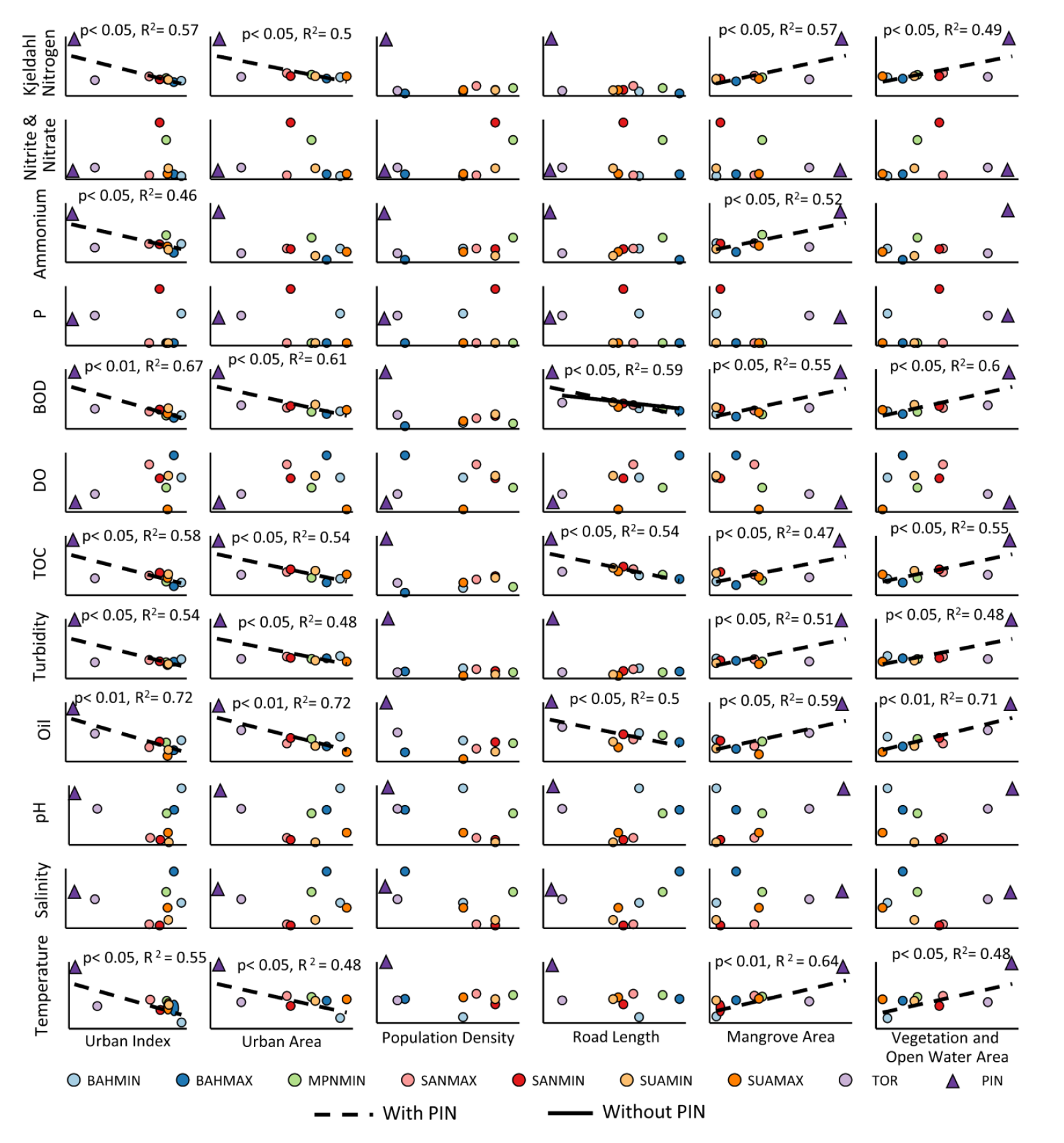
Scatter plots and linear models of surface water chemistry metrics as predicted by surrounding land cover. Lines show statistically significant models (p <0 .05). Dashed lines are those with all sites and the solid line is that without PIN, which was an outlier and responsible all but one of the significant models. Biochemical oxygen demand, nitrogen concentrations, organic carbon, turbidity, oil and grease, and temperature are all predicted to decrease with increasing urbanization.

Flooding metrics resulted in much fewer significant linear models compared to surface water chemical properties when predicted by surrounding land coverage (Figure 8). In t-tests between urban and non-urban sites, however, in which urban is defined as any urban cover within the 0.5 km sampling radius, most metrics resulted in significant differences. Mean water depth and the proportion of time flooded were the only flooding metrics to result in significant linear models (p < 0.05). In both cases, the surrounding open water and vegetation coverage was the strongest predictor and resulted in positive linear relationships. However, much of the significance of these models may only be due to the highly urban outlier site of MPNMAX, and none of the models resulted in significant slopes when this site was removed from the analysis. In binomial comparisons of urban and non-urban sites through t-tests, all metrics except flooding frequency were statistically different. These tests suggest urban sites have lower depth but longer hydroperiod and overall lower proportion of time flooded.

**Figure 8.**
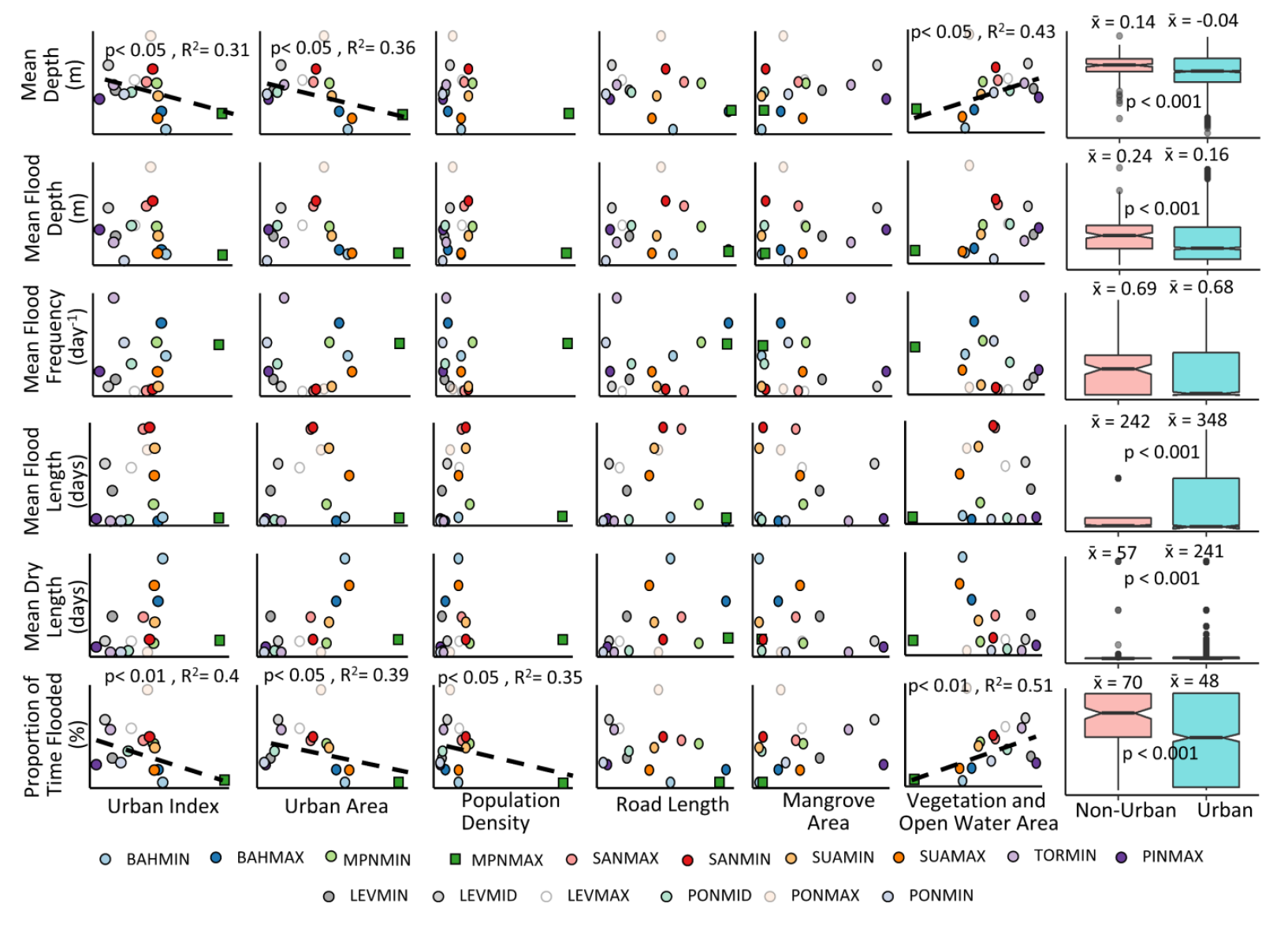
Scatter plots, linear models, and boxplots of flooding metrics as predicted by surrounding land cover. In scatterplots, lines show statistically significant models (p <0 .05). None of the linear models could be repeated without the outlier of MPNMIN. In boxplots, Urban is defined as those points with any urban land cover within the 0.5 km sampling radius. P-values are the result of student t tests and are displayed only when a significant difference resulted between urban and non-urban values. Under this definition, there are significant differences in urban and non-urban mangroves in all flooding metrics except flood frequency. These results suggest urban mangrove are shallower and flooded less of the time than non-urban mangroves.

In testing the response of specific tidal and rainfall constituents to surrounding land coverage, there were mixed results. The amplitudes of two tidal constituents were most explained by the surrounding urban index, in which a negative logarithmic relationship was modeled (Figure 9a), although there was no relationship between the combined amplitude of all constituents and surrounding land cover. While these models suggest lower contributions from some tidal constituents in urban mangroves, the contribution from rainfall was not found to be dependent upon urbanness, and instead geomorphology and tidal connectivity were more important (Figure 9b). In this case, partially restricted hydro-geomorphologies responded by increasing water levels five times that of cumulative rainfall. This was significantly greater than the response by open systems at 0.5 times cumulative rainfall (t test; difference = 4.2 cm/cm, p < 0.05). There was no relationship between the extent of water level response to rainfall and surrounding urban coverage across sites (p > 0.5).

**Figure 9.**
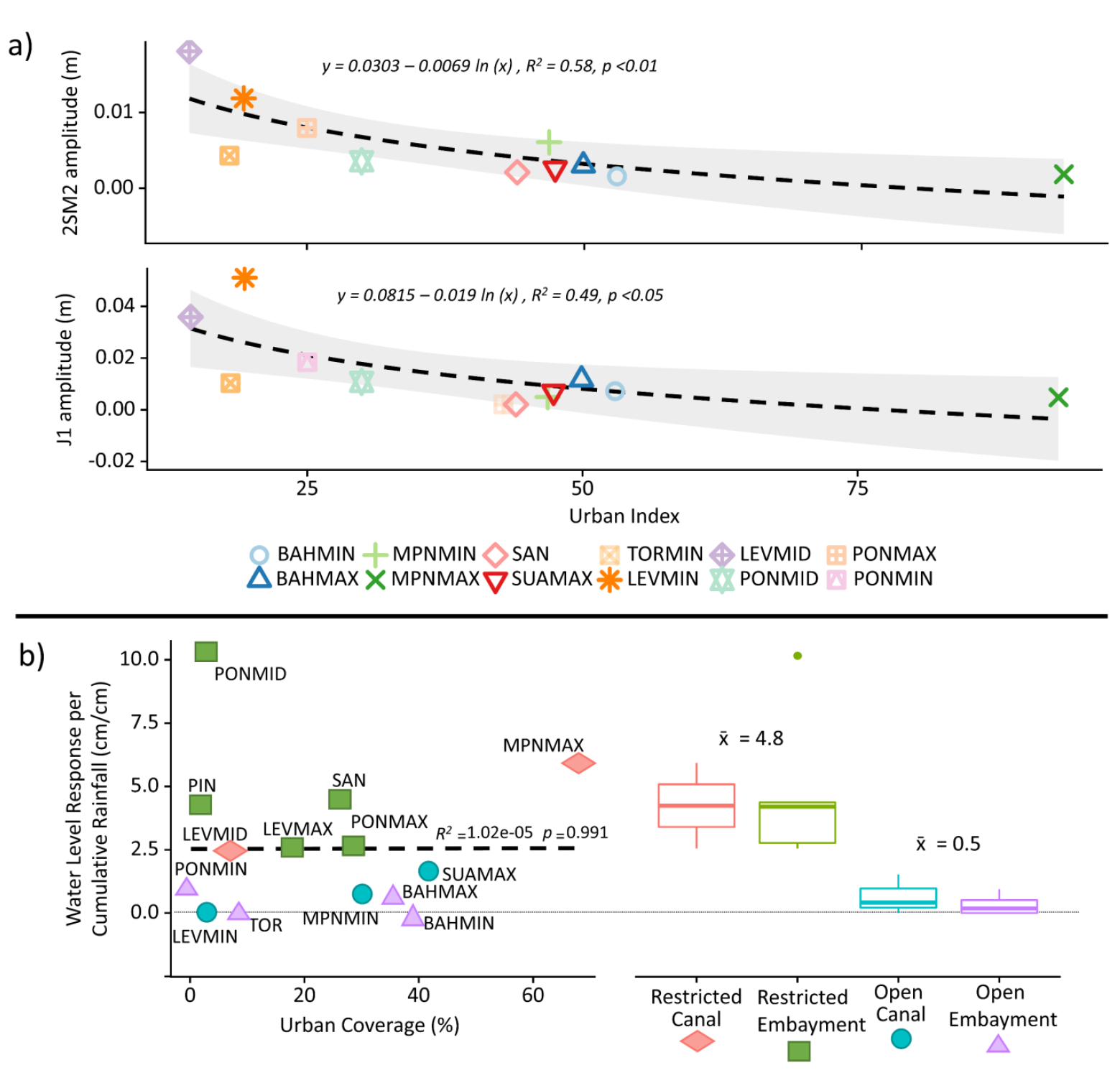
Mixed evidence for the influence of urban land use on mangrove hydrology. The shallow water semidiurnal (2SM2) and the smaller lunar elliptical diurnal (J1) tidal constituents show decreasing amplitude with increasing urban coverage (a). But there was no evidence for an effect of urbanization on water level responses to rainfall (b). Instead, differences in rainfall response were best explained by geomorphology, where partially restricted waterbodies respond more sharply to rainfall than to those that are more open to tidal connections.

## DISCUSSION

There was mixed evidence for a significant influence of urban land cover on flooding dynamics or surface water chemical properties across the mangrove study sites. Both mean depth and proportion of time flooded were best modeled to increase with surrounding open water and vegetation coverage and decrease with surrounding urbanness (Figure 8). Yet the models presented in Figure 8 were no longer significant when MPNMAX was removed from the analysis, suggesting the site may be an unrepresentative outlier. Still, such a result would be expected from a decrease in tidal amplitudes in urban sites, which is consistent with indirect findings from another study (Marois and Mitsch 2017). Binomial comparisons of urban vs. non-urban corroborated this by showing lower average depths and flooding times, but longer hydroperiods in urban sites (Figure 8). This implies urban sites do not flood as often, but when they do, flooding is shallower and more prolonged. This is further corroborated by analyses on individual constituents, in which the amplitudes of the J1 (smaller lunar elliptical diurnal) and 2SM2 (shallow water semidiurnal) constituents were found to decrease as urbanization increased (Figure 9a). The difference in amplitudes between the least and most urban sites is around 3 and 1 cm, respectively, for the J1 and 2SM2 constituents, which may be enough to lead to lower average depths in some urban forests.

The potential explanation for a decrease in tidal amplitudes with urbanization is unclear. It’s possible this is in response to urban storm water infrastructure, which may exacerbate rainfall runoff to the point in which it interferes with otherwise normal tidal forcing. This would be consistent with urban hydrology theory (Leopold 1968; Hollis 1975; Lee et al. 2006; Dietz and Clausen 2008). But tidal models were constructed in the absence of rainfall within a 4-day period, so all tidal amplitudes should be independent of the influence of precipitation. Further, there was no significant effect of urbanization on the water level responses to rainfall (Figure 9b). Instead, tidal connectivity resulting from geomorphology was found to be more important, where restricted and less connected systems respond more strongly to rainfall than do open and more connected systems (Figure 9b). Urbanization might therefore be influencing tidal components through engineered changes to geomorphology.

Canalization and dredging, for example, are sometimes associated with highly urban landscapes, and are also likely influencing tidal amplitudes through changes in geomorphology. There was some evidence for this in the results. Water levels in the dredged portion of Martín Peña canal (MPNMIN) are nearly identical to those at the mouth of the Río Puerto Nuevo in the San Juan Bay (BAHMAX) (Figure 3). Farther into the un-dredged and highly constricted Martín Peña (MPNMAX), however, daily water ranges are nearly half that of MPNMIN and the influence of rain on the water level model is five times greater (Table 4, Figure 9b). Thus, it is likely that the dredging of the lower portions of the canal has allowed greater tidal influence and limited response to rainfall. A similar reduction in tidal amplitude and amplification of the importance of rain is seen between SUAMAX and SUAMIN (Figure 3). These two portions of the Suarez canal are separated by the Baldorioty expressway (PR highway 66) and a network of frontage roads that form a highly engineered constriction, reducing the canal width from roughly 50 to 15 meters. The restricted portion of the canal at SUAMIN and San Jose Lagoon stations (SANMIN and SANMAX), the daily range in water levels is 3.5 times less than that at the tidally connected SUAMAX and the importance of rain in the water level model is nearly six times greater (Table 4, Figure 9b). Thus, the constriction of the canal at the highway crossing is likely impeding flood drainage, resulting in greater importance of rainfall and limited tidal connectivity in the upper half of the canal.

With surface water chemical properties, the most significant models could not be repeated without the extreme values of Piñones lagoon (PIN), which is a relatively pristine system with a weak hydrologic connection to the rest of the estuary (Ellis 1976; Lugo et al. 2011). Biological oxygen demand, however, was modeled to decrease as surrounding road length increased in models with and without PIN. But the effect of street lengths on BOD is opposite of that from previous studies (Mallin et al. 2009; Erickson et al. 2013), which have shown urban storm water runoff to increase BOD by introducing oxygen demanding substances. The observed trend from Figure 7 is thus probably spurious as it is unlikely street length in San Juan has the opposite effect on BOD as other aquatic systems. Instead, the observed trend is likely due to elevated organic carbon (TOC) and nitrogen (Kjeldahl nitrogen) at Piñones (Sawyer 2003). A linear model constructed of these two predictors performed far better than any of the tested land cover metrics (BOD ∼TOC + Kjeldahl nitrogen; R^2^ = 0.98, p < 0.01). Apart from BOD, TOC and Kjeldahl nitrogen, Piñones held significantly higher ammonia, turbidity, oil and grease, and temperature, than most other sites, and all trends suggest the opposite of previous studies in which the least urban water bodies hold the lowest nutrient and contaminant concentrations (McClelland and Valiela 1998; Mallin et al. 2009; Erickson et al. 2013). Thus, Piñones being a relatively pristine system, it must be inferred that its anomalous surface water chemistry is explained more by its minimal tidal influence than by urban inputs. This further demonstrates that hydrology and water chemistry must by studied within the context of geomorphology and tidal connectivity when assessing anthropogenic influences (Kjerfve et al. 1999; Adame et al. 2010).

There are other examples of isolated abnormalities that could be associated with either urbanization or natural variations in geomorphology and water chemistry. For the former, nitrogen was abnormally high in various forms in some of the water bodies of the San Juan Bay Estuary, providing further evidence for the commonly reported eutrophication of the system (Webb and Gómez-Gómez 1998; Cerco et al. 2003). Ammonia on average was four times higher in Piñones, a relatively pristine site, than the other water bodies, and total Kjeldahl nitrogen was ten times higher. Similar patterns have been seen in other landward mangroves with less tidal connectivity and some human inputs (Tam and Wong 1998). SANMIN, a moderately urban site, also registered higher nitrate and nitrite concentrations than all other water bodies, this time likely due to an excessive amount of wastewater entering the lagoon from the surrounding area through the San Antón creek (Gómez-Gómez and Quiñones 1983). Sewage may also be to blame for high phosphorous concentrations at BAHMIN, which includes the mouth of Malaria canal and the intermittent effluent of sewage (Cruden 2015). Piñones was again singled out from the other water bodies in its flooding dynamics, along with LEVMAX, PONMAX, and PONMID, all of which exhibited significantly greater hydroperiods and maximum flood depths in comparison with the other sites. Again, the likely reasons for these anomalies are mixed. These mangroves are characterized by relatively isolated inland lagoons with little tidal connectivity. As a result, they are more sensitive to precipitation inputs (Rodríguez-Martínez and Soler-López 2014), which induce rapid and prolonged flooding. At PONMAX and LEVMAX, the isolation is likely by design, with both serving as urban storm water retention basins. But Piñones and PONMID are both protected and relatively undisturbed sites with minimal human influence, thus their abnormal hydroperiods are more likely a result of natural forces.

This study used water level models based on months and in some cases years of observations across sixteen sites in Puerto Rico. Most of the models explained more than 50% of the observed variations in water levels, and many of them explained at least 70% of the variations. Still, unexplained variations may be a result of untested variables or biased or inaccurate results. One potential source of error is the vertical accuracies of the DEMs (Table 1), which although well within the range of most water levels, are still greater than some of the observed differences in depths between sites. More accurate DEMs that cover such a large area may be available for future studies and could be used to improve upon this study. Other improvements can be made by capturing more long-term observations across similarly quantified urban gradients, and by capturing the response to rainfall, which was the primary source of unexplained variation. These studies should consider land cover alongside geomorphology and tidal connectivity, and should attempt to isolate tidal constituents and specific urban components. They should also use varying definitions of urbanness, as it was demonstrated here that binomial classifications of urban and non-urban resulted in significant differences while gradient analyses did not. Doing so will identify potential causal factors and will be an increasingly important task for mangrove management in the Anthropocene.

As urbanization continues to drive ecosystem processes and habitat loss, its influence on mangrove hydrology is not yet well understood. While changes to hydrology and water chemistry are well documented in urban systems, there remains little empirical evidence for specific influences along quantified urban gradients in mangrove systems. This study shows that such evidence remains confounded with other important factors along three urban gradients in Puerto Rico. Although there was some evidence for changes in tidal amplitudes and surface water chemical properties in urban sites, there were few consistent trends to tie any variations in water levels or chemistry to surrounding urbanization. Geomorphology and tidal connectivity were found to be important influences, both of which exhibit both natural and anthropogenically induced variations. Future studies must therefore distinguish between specific components of urban landscapes and natural or engineered variations in geomorphology. They may also improve upon the results presented here by using more accurate DEMs and by better capturing the uncertainty in water level responses to rainfall.

## ACKNOWLEDGEMENTS

Research was partly funded by the United States Forest Service International Institute of Tropical Forestry, San Juan, Puerto Rico. Ariel Lugo, Ferdinand Quiñones, and Autumn Oczkowski provided feedback on first drafts.

